# Predicting the similarity of two mass spectrometry runs using only MS1 data

**DOI:** 10.1101/2023.11.29.569301

**Authors:** Abdullah Shouaib, Andy Lin

**Author notes:** Corresponding author: 1100 Dexter Ave N, Suite 500, Seattle, WA 98109.

## Abstract

**Background:** Traditionally researchers can compare the similarity between a pair of mass spectrometry-based proteomics samples by comparing the lists of detected peptides that result from database searching or spectral library searching. Unfortunately, this strategy requires having substantial knowledge of the sample and parameterization of the peptide detection step. Therefore, new methods are needed that can rapidly compare proteomics samples against each other without extensive knowledge of the sample.

**Results:** We present a set of neural network architectures that predict the proportion of confidently detected peptides in common between two proteomics runs using solely MS1 information as input. Specifically, when compared to several baseline models, we found that the convolutional and siamese neural networks obtained the best performance. In addition, we demonstrate that unsupervised clustering techniques can leverage the predicted output from our method to perform sample-level characterizations. Our methodology allows for the rapid comparison and characterization of proteomics samples sourced from various different acquisition methods, organisms, and instrument types.

**Conclusions:** We find that machine learning models, using only MS1 information, can be used to predict the similarity between liquid chromatography-tandem mass spectrometry proteomics runs.

## 1 Background

There is a plethora of historical liquid chromatography-tandem mass spectrometry (LC-MS/MS) proteomics data that have been deposited into online repositories such as PRIDE [1] and MassIVE [2]. When presented with new samples, researchers may want to characterize these samples by comparing them to existing historical data. Performing this analysis requires the ability to measure the similarity between proteomics runs. Unfortunately, sample-level comparison methods for mass spectrometry (MS) runs are scarce due to the difficulty of measuring the similarity between a pair of runs. This difficulty is due to data differences that result from biologically irrelevant factors such as instrument acquisition methods, instrument type, and sample preparation protocol.

Previously developed methods for measuring the similarity between a pair of mass spectrometry-based proteomics runs include counting the number of confidently detected peptides in common [3] and counting the fraction of pairs of fragment spectra (MS2), out of all pairs, that have a spectra dot product above a threshold [4, 5, 6]. Though successful, these approaches are not trivial given that they require a peptide detection step (i.e., database or spectral library search). This peptide detection step is time consuming, resource intensive, and requires a great deal of parameterization. In addition, MS2 spectra that are collected by data-dependent acquisition (DDA) and data-independent acquisition (DIA) are fundamental different from one another, and these aforementioned methods do not account for these differences.

To circumvent the limitations of these MS2-based methods, a method called MS1Connect was developed to measure the similarity between pairs of mass spectrometry runs using only intact peptide (MS1) data [7]. However, the MS1Connect objective function was handcrafted and therefore may not be ideal. In addition, this method can be computationally expensive since it requires solving a maximum bipartite matching problem for each pair of runs.

Rapid advances in the machine learning (ML) field now enable ML algorithms to learn scoring functions in a broad range of applications, supplanting the need for handcrafted metrics. For example, MS2DeepScore uses a Siamese neural network to directly predict the structural similarity of a pair of metabolites from their MS2 spectra [8]. In proteomics applications, neural network architectures like GLEAMS [9] and DLEAMSE [10] have also been developed to generate embeddings of MS2 spectra for the downstream purpose of clustering and analysis. While successful, these methods ultimately measure the similarity between pairs of spectra and not runs, which can contain tens of thousands of MS2 scans.

In this work, we aimed to determine whether machine learning methods can directly predict the similarity between a pair of proteomics runs using only MS1 information as input. Previous work has shown that MS1 information is informative and can be used as input into classification tasks. For example, MS1 features have been successfully used for classifying between tumor and normal tissue samples [11, 12]. In addition, as noted above, MS1 features has been successfully used to measure the similarity of a pair of runs [7]. Therefore we hypothesized that it would be possible to use machine learning to directly predict the similarity between a pair of proteomics-based mass spectrometry runs using only MS1 features.

Here, we present a convolutional neural network architecture that takes as input the MS1 features from two different runs, and predicts the pairwise similarity between them. Specifically, our model predicts the Jaccard index between the sets of confidently detected peptides of two runs. To our knowledge, this is the first method that predicts the similarity between a pair of runs using solely MS1 data as input, meaning that peptide detection step was not performed.

We provide evidence that our model successfully measures the similarity between two proteomics runs by comparing the predicted value to the ground truth value. Our approach outperforms a non-machine learning baseline method and other machine learning models, such as a random forest (RF) and support vector machines (SVM). In addition, we demonstrate that unsupervised clustering of the affinity matrix generated by the predicted scores may have utility in characterizing unlabelled samples. In this case, we show that clustering the samples using the predicted scores can be used to suggest metadata labels such as species type. Finally, we observe that our method can potentially be applicable to samples that originate from organisms that fall outside the training dataset.

## 2 Methods

### 2.1 Training and test dataset

A total of 966 RAW files, each from a unique PRIDE ID, were downloaded from PRIDE [1] using the ppx package (version 1.2.4) [13]. Every RAW file was converted to mzML format using ThermoRawFileParser (version 1.2.0) [14]. These runs originated from six different species (Table 1), and the list of filenames and PRIDE ID’s acquired for this study can be found in the Supplemental File 1.

**Table 1:** Organism percent coverage. The number and percentage (in separateness) of files in the training and test set. A roughly equal percentage between the training and external test set informs us that the datasets are fairly balanced in regards to organism classifications of the samples.

| Organism | Train | Test |
| --- | --- | --- |
| *Homo sapiens* (human) | 634 (70.4%) | 43 (65.2%) |
| *Mus musculus* (mouse) | 69 (7.67%) | 7 (10.6%) |
| *Escherichia coli* | 67 (7.44%) | 5 (7.58%) |
| *Drosophila melanogaster* (fruit fly) | 50 (5.56%) | 5 (7.58%) |
| *Arabidopsis thaliana* (mouse-ear cress) | 47 (5.22%) | 4 (6.06%) |
| *Saccharomyces cerevisiae* (baker’s yeast) | 33 (3.67%) | 2 (3.03%) |

Of the 966 runs, 900 runs were used to generate the training set while the remaining 66 were used to generate the test set. The 900 runs yielded 404,550 unique training pairs, while the test runs yielded 2,145 pairs. To both minimize and assess overfitting, the training set was split into a training and validation set via an 80/20 split on pairs. As a result, a validation set of 80,910 pairs was generated.

To assess whether our model could predict the pairwise similarities of proteomics runs from organisms that fall outside the training set, we also evaluated a set of five zebrafish and five hamster runs. These 10 runs were also compared to five human runs from the original training dataset. Details on these datasets can be found in Supplemental File 2. Note that the five hamster runs originated from two different species: *Mesocricetus auratus* and *Cricetulus griseus*.

### 2.2 Representation of Mass Spectrometry Runs

We represent each run as a matrix of discretized MS1 features, where rows represent retention time bins, columns represent *m/z* bins, and each cell contains the summated intensity within that bin. To obtain the MS1 features, we performed MS1 feature detection using PyOpenMS (version 2.7.0) [15]. Each MS1 feature is represented by the following three values: mass over charge ratio (m/z), retention time (RT), and intensity. The retention times were normalized on a scale of zero to one based on the proportion of total ion current (TIC) detected up to a given time. We then removed MS1 features with a normalized retention time ≤ 0.05 and ≥ 0.95. In addition, the intensities were normalized to a range from zero to one by dividing by each intensity by the maximum intensity among all MS1 features in each run. Finally, the MS1 features were discretized by binning the *m/z*, with a width of 1.0005079 Da, and normalized retention time, into one, two, or five bins.

### 2.3 Database Searching

For our ground truth values we calculated the Jaccard index between the two sets of confidently detected peptides after performing a database search on two runs. For each run we performed the database search using the Comet search engine [16] within the Crux toolkit (version 4.1) [17, 18]. Spectra were searched against the Uniprot Swiss-Prot database that was downloaded in January 2022 [19]. The database search parameters were generated using Param-medic [16, 20]. Following the search, Percolator was used to the improve the number of detections and estimate the false discovery rate [21]. The resulting detections were filtered down to a 1% FDR at the PSM level.

Once the set of confident PSM detections for each run was determined, we calculated the pairwise similarity between any two runs using the Jaccard index. We defined the Jaccard index as

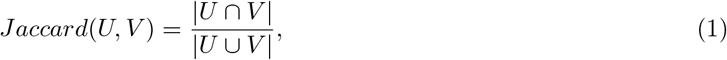

where *U* and *V* are the set of detected peptides, at a 1% FDR threshold, from each run respectively.

### 2.4 Prediction of pairwise similarity score

We tested five different machine learning methods and one non-ML baseline for predicting the similarity between pairs of runs. Each method takes as input the matrix representation of two runs and outputs the predicted Jaccard index of the confidently detected peptides. We evaluated the performance of each method using the mean squared error (MSE) loss function between the predicted and database-search derived Jaccard coefficients. In addition to MSE, we also calculated the root mean squared deviation (RMSE), mean absolute error (MAE), and Pearson correlation coefficient.

#### 2.4.1 Spectral Dot Product Approach

To ensure our machine learning approaches obtained reasonable results, we compared them to the spectral dot product baseline. Specifically, we measured the similarity between a pair of proteomics runs by calculating the dot product between the discretized MS1 features.

#### 2.4.2 Support Vector Regression and Random Forest Models

We first trained several support vector regression (SVR) and random forest (RF) models on the task of predicting Jaccard indices using a matrix representation of MS1 features as input. A total of three SVR and three RF models were trained, where in each instance we set the number of retention time bins to one, two, or five. The SVR models were trained with a regularization parameter of 1.0 using the radial basis function kernel. The RF models were trained with 100 decision trees, and the MSE was set as the criterion to measure the quality of the split.

#### 2.4.3 Deep Neural Network Architectures

In addition to the SVR and RF models, we trained two different neural networks. The first neural network, which we refer to as a deep neural network (DNN) in this manuscript, contains an input layer, three fully connected layers (with unit sizes of 512, 128, and 32 respectively), two dropout layers with a rate of 0.2, and a final output layer.

The second neural network we trained was a 2-dimensional convolutional neural network (CNN). The CNN model takes as input the matrix representations of two proteomics runs simultaneously by vertical stacking the matrix form of the two runs. The architecture of this network consisted of two convolutional blocks followed by a fully connected layer and then a output layer. Each block contained two convolutional layers, a max pooling layer, and a dropout layer. The convolution window size and stride length was set to (2,6) and two. In addition, the max pooling layer had a pool size and stride length of (2,2) and two. The output of the convolution blocks are passed to a L2-regularized, 32-dimensional fully connected layer, and then to the L2-regularized output layer.

We trained the DNN for 30 epochs and trained the CNN for 90 epochs. Both architectures used a learning rate of 0.0002. All network layers use the scaled exponential linear units (SELU) activation function [22]. The fully connected layers were initialized using LeCun normal initialization and the convolutional layers were initialized using the Glorot uniform initialization [23, 24]. In addition, each iteration consisted of a batch size of 512. Training and evaluation were performed on Intel Xeon Gold 6230 processors with 376 GB of memory and an NVIDIA GeForce RTX 2080 Ti graphics card.

#### 2.4.4 Siamese Neural Network Architecture

In addition to the CNN and DNN, we also trained a siamese neural network architecture to perform the same task (Figure 1). A siamese network contains two identical subnetworks where weights are tied to each other across the subnetworks. The MS1 binned intensities of a single proteomics run is passed as input into each subnetwork. Each subnetwork contains two convolutional blocks, a fully connected layer, and a embedding layer. Each convolutional block contained two one-dimensional convolutional layers, a max-pooling layer, and a dropout layer. The convolution layers had a window length of 3 and a stride length of 1. In addition, the max pooling layers consisted of a pool size of 1 and stride length of 2. Finally, 20% of nodes were randomly dropped in the dropout layer.

**Figure 1:**
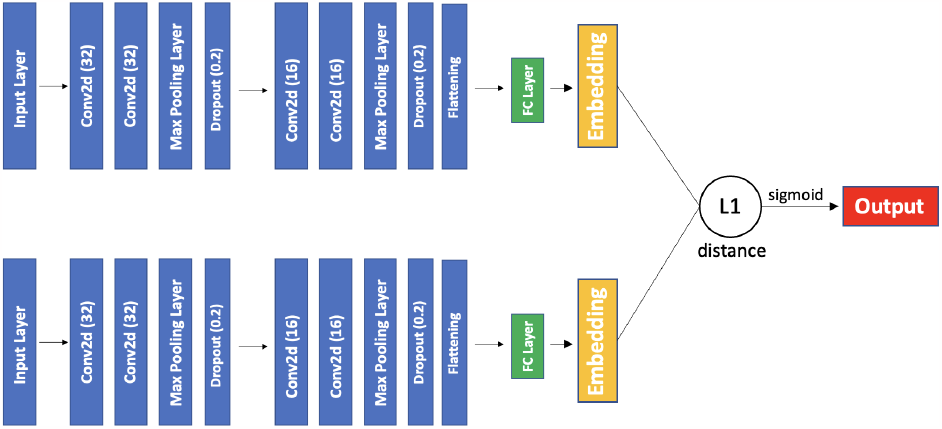
Siamese model architecture. The Siamese network contains two instances of the same convolution blocks and fully connected layers. The filter sizes of the convolution layers are denoted in parenthesis, while the rate of dropout is denoted in the dropout layers. The discretized MS1 features of sample 1 are passed through one network to create an embedding, while sample 2 passes through the other to create another embedding. The L1 distance between the two embeddings is then calculated as the output.

The output of each subnetwork is passed to a final, L2-regularized, 32-dimensional fully connected layer to create the final embedding layer. The output of the Siamese network is the L1 distance between the two embeddings. The Siamese neural network was trained for 40 epochs using the Adam optimizer with learning rate 0.0002. Each iteration consisted of a batch size of 256.

Like the CNN and DNN, all network layers used the scaled exponential linear units (SELU) activation function [22]. The fully connected layers were initialized using LeCun normal initialization and the convolutional layers were initialized using the Glorot uniform initialization [23, 24].

### 2.5 Code Availability

All code was developed using Python 3.10. PyOpenMS (v 2.7.0) was used to process all MS runs and detect MS1 features. The RF and SVR models were implemented using Scikit-Learn (v 1.0.1), while the neural networks were implemented using the Tensorflow/Keras framework (v 2.4.0). Hierarchical clustering and t-SNE were performed using Scikit-learn, SciPy (v 1.8.1), and Seaborn (v 0.11.2).

## 3 Results

### 3.1 Neural networks outperform other models for similarity learning using only MS1 features

To determine whether our machine learning methodology can reliably predict the similarity between a pair of mass spectrometry-based proteomics runs, we compared the predicted Jaccard index against the ground truth Jaccard index. Specifically, we measured the MSE for the training, validation, and test set. In addition to MSE, we also calculated additional metrics such as RMSE, MAE, and Pearson correlation.

Our results showed that the CNN and two different random forest models obtained the best performance on the test set with a MSE of 0.005 (Table 2). On the test set, the CNN also obtained the best performance, as measured by MAE and Pearson correlation, while the two random forest models obtained the lowest RMSE. We found that the CNN and the third random forest model both obtained similar middling performance on the test set. In addition, in general, we found that the three SVR models and the DNN model obtained the worst performance on the test set.

**Table 2:** Performance obtained by each model. A table of the performance obtained by each model, as measured by mean square error (MSE), root-mean-square error (RMSE), mean average error (MAE), and Pearson correlation (Pearson), on the training and test set. Bold values indicate the best value for each column.

| Model | # RT bins | Training Set |  |  |  | Validation Set |  |  |  | External Test Set |  |  |  |
| --- | --- | --- | --- | --- | --- | --- | --- | --- | --- | --- | --- | --- | --- |
|  |  | MSE | RMSE | MAE | Pearson | MSE | RMSE | MAE | Pearson | MSE | RMSE | MAE | Pearson |
| SVR | 1 | 0.00476 | 0.069 | 0.066 | 0.50 | 0.0048 | 0.069 | 0.066 | 0.50 | 0.0045 | 0.067 | 0.064 | 0.32 |
| SVR | 2 | 0.00490 | 0.070 | 0.067 | 0.48 | 0.0049 | 0.070 | 0.067 | 0.47 | 0.0049 | 0.070 | 0.067 | 0.28 |
| SVR | 5 | 0.00510 | 0.071 | 0.068 | 0.42 | 0.0051 | 0.071 | 0.068 | 0.41 | 0.0053 | 0.073 | 0.070 | 0.19 |
| RF | 1 | **0.00003** | **0.005** | **0.003** | **0.99** | 0.0002 | 0.014 | 0.009 | 0.89 | **0.0005** | **0.022** | 0.017 | 0.56 |
| RF | 2 | **0.00003** | **0.005** | **0.003** | **0.99** | 0.0002 | 0.014 | **0.007** | 0.91 | **0.0005** | **0.022** | 0.017 | 0.55 |
| RF | 5 | **0.00003** | **0.005** | **0.003** | **0.99** | 0.0002 | 0.014 | 0.008 | 0.90 | 0.0007 | 0.026 | 0.018 | 0.41 |
| DNN | 1 | 0.00060 | 0.025 | 0.017 | 0.60 | 0.0004 | 0.019 | 0.017 | 0.60 | 0.0015 | 0.039 | 0.029 | 0.11 |
| CNN | 1 | 0.00020 | 0.014 | 0.009 | 0.91 | 0.0002 | 0.015 | 0.010 | 0.88 | **0.0005** | 0.023 | **0.015** | **0.59** |
| Siamese | 1 | 0.00010 | 0.009 | 0.007 | 0.96 | **0.0001** | **0.001** | **0.007** | **0.95** | 0.0008 | 0.028 | 0.021 | 0.40 |

While the CNN and RF obtained the best performance on the external test set, we found that the siamese network obtained the best performance on the validation set. For the training set, we found that all three random forest models, at varying retention time bins, obtained the best performance, as measured by all four metrics. Specifically, these models obtained a performance of 0.00003, 0.005, 0.003, and 0.99 for the MSE, RMSE, MAE, and Pearson correlation, respectively. Given that the siamese is best performing on the validation set, and given its performance on the training set as well, it is understood that the model is overfitting to the MS runs in the training set.

Focusing on the SVR and RF models, we found that, in general, the number of retention time bins did not significantly change performance. Specifically, we found that the MSE, RMSE, and MAE did not significantly change as a function of the number of retention time bins. In addition, we found that the Pearson correlation largely did not change as a function of the number of retention time bin for the random forest models.

In regards to the support vector regression models, we found that the Pearson correlation decreased as a function of the number of retention time bins. This decrease was observed for the training, validation, and test set. For example, the Pearson correlation of the test set decreased from 0.32 to 0.19. as the number of retention time bins increased from one to five. We speculate that this decrease in performance could be explained by edge effects created by binning retention time.

We observed that both our Siamese network and the CNN were learning, given the decrease in the loss value over epochs (Figure 2). This observed decrease in error demonstrates that neural network architectures can learn some relationship between MS1 features and MS2-based attributes. On the other hand, we observed that the DNN did not learn enough during the training regime (Figure S2) This could be due to size of the DNN, which may not be sufficiently complex enough to capture the intricate relationships in the data. Another reason this could be the case is that CNNs take into account the spatial and sequential structure of the data, making them potentially better suited for mass spectral data compared to artificial neural networks. To ensure our models obtained reasonable performance, we also compared their performance to the performance obtained by the spectral dot product score. We found that the spectral dot product score obtained poor performance. This score obtained a MSE and Pearson correlation of 0.0715 and 0.18, respectively, on the test set. In addition, this score obtained a MSE and Pearson correlation of 0.0877 and 0.30, respectively, on the validation set.

**Figure 2:**
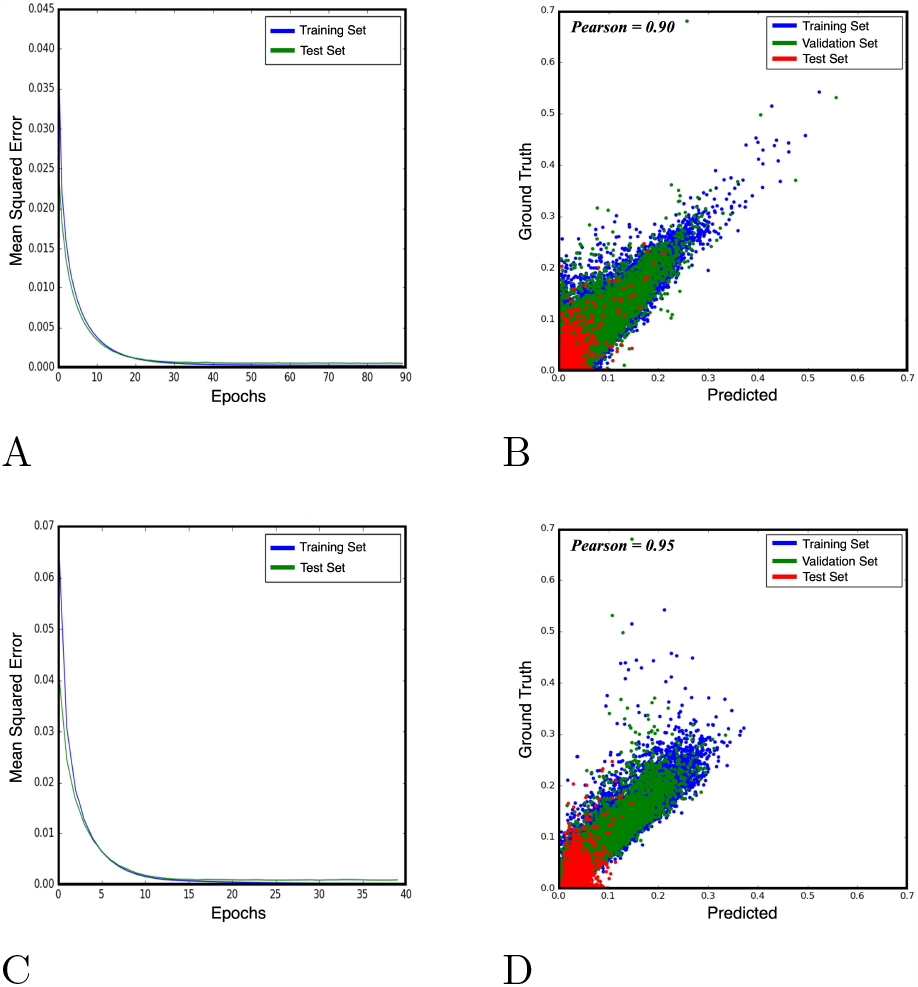
Loss curve and correlation of predicted and ground truth scores for CNN and Siamese architectures. (A) MSE loss curve for the CNN model. (B) A scatter plot of the predicted score versus the ground truth scores for the CNN model. (C) MSE loss curve for the Siamese network. (D) A scatter plot of the predicted score versus the ground truth scores for the siamese model. Together these plots indicate that CNN and siamese models can learn from MS1 feature representations.

### 3.2 Characterizing MS samples through unsupervised clustering of predicted similarities

After determining that some neural network architectures can successfully predict the Jaccard index of detected peptides using solely MS1 features, we then investigated whether unsupervised clustering of these predicted pairwise similarity scores could aid in sample comparison and characterization. Unsupervised clustering for sample characterization has been successfully used in a variety of disciplines ranging from metabolomics to proteomics [6, 9, 8]. To test our hypothesis, we preformed hierarchical clustering (HCA) and t-Stochastic Neighbor Embedding (t-SNE) on a similarity matrix generated from our Siamese model for the training dataset.

After performing dimensionality reduction using t-SNE on the predicted similarities, we found that the presence of several clusters that corresponded with the organism a sample originated from (Figure 3). For example, a majority of runs from *Drosophila melanogaster* (fruit fly), *Arabidopsis thaliana* (mouse-ear cress), and *E. coli* cluster into distinct groups. On the other hand, we found that many of the mouse runs overlap with the human runs, indicating a higher similarity among these samples. We speculate this occurs because of the high homology between the two species. In addition, we note that there are many instances where points from different species lie close to one another. These occurrences could be explained by other relevant differences between samples such as tissue type, instrument, modifications, incorrect metadata, etc. Finally, we obtained similar results when we performed a hierarchical clustering on the data (Supplemental Figure 1)

**Figure 3:**
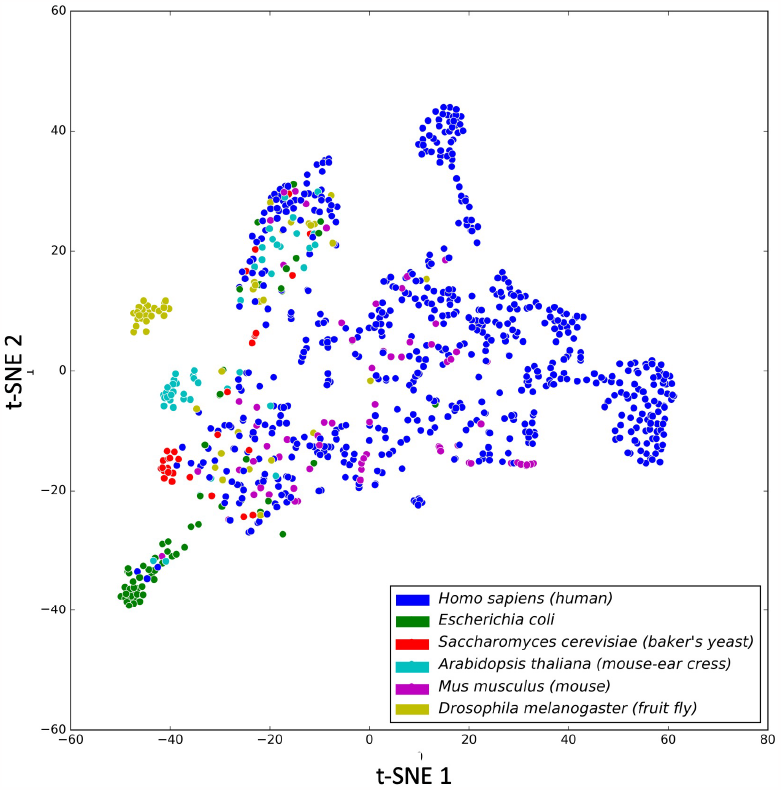
t-Stochastic Neighbor Embedding (t-SNE) applied to the similarity matrix from the training dataset. Each point represents an individual run and is colored based on organism label.

### 3.3 Characterizing samples from species outside the training set

Machine learning models are beholden to the limitations of their training data. It is understood that novel datasets can be acquired which fall outside the domain of the training data, and in such cases the model may not be applicable to those outliers. This drawback applied to our model since it was only trained on data from a subset of model organisms. To test whether our predictive model could be generalizable enough to predict similarities for organisms not in the training dataset, we evaluated the performance of our model on runs from two species not found in the training set: zebrafish and hamster.

Of each outlier organism, five MS runs were acquired and discretized through our siamese model to predict the pairwise Jaccard similarities. We then compared these runs to five randomly chosen human samples from the original training dataset as a point of reference. The Pearson coefficient between the ground truth and predicted Jaccard similarities were found to have a moderate correlation of 0.60 (Figure 4), while the MSE was 0.0009, and the MAE was 0.0208. This correlation suggests our siamese model can be potentially applicable to MS runs from organisms outside the training set, though much more data from more diverse organisms aren’t observed here.

**Figure 4:**
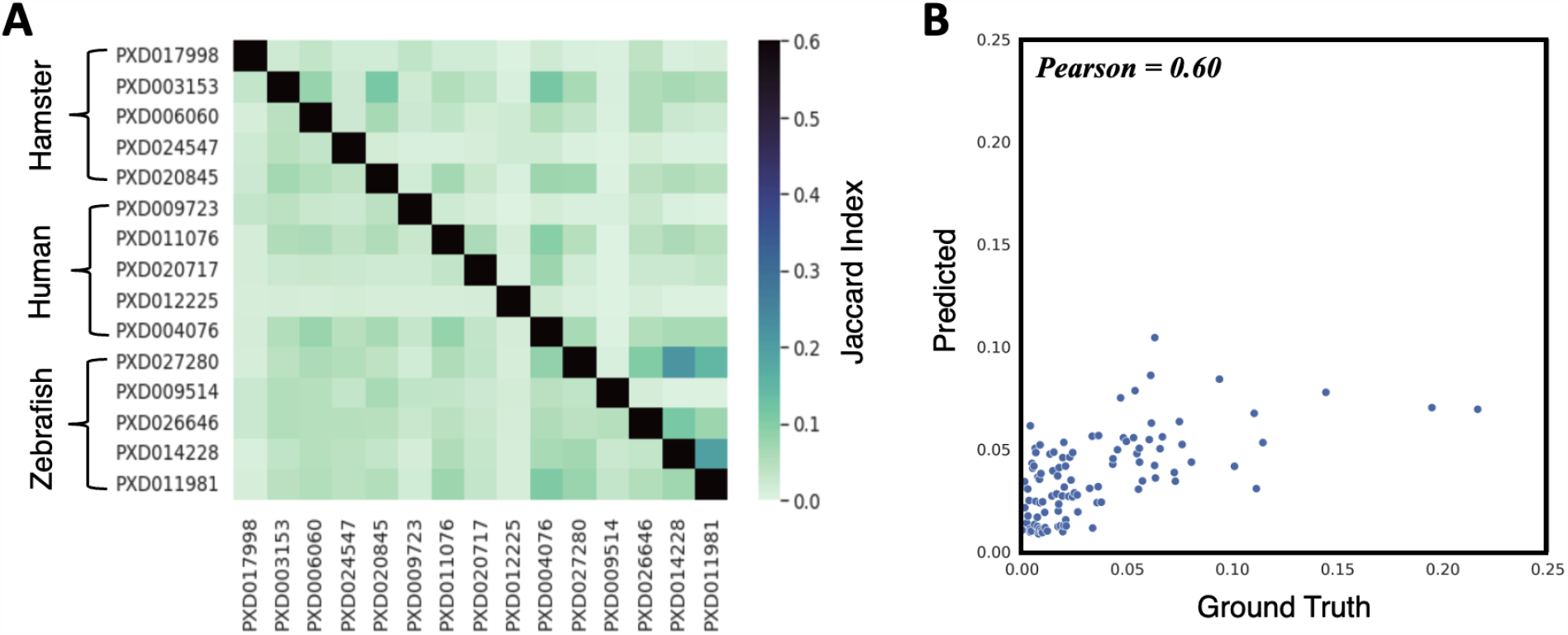
Predicting similarities of species not in training set. (A) A heatmap of the predicted and ground truth Jaccard similarity values between five hamster, human, and zebrafish runs. The upper triangle of the heatmap contains ground truth values while the lower triangle represents the predicted values. The black diagonal line across the diagonal does not represent anything as we did not predict or calculate self-similarity. (B) A scatterplot view of the data found in (A). The Pearson correlation between the ground truth and predicted scores for the unique pairs in the external and humans samples is 0.60.

## 4 Discussion

In this work, we investigate several machine learning models to rapidly predict the similarity between different proteomics runs at the sample level. Specifically, these models use solely MS1 features as input to predict the pairwise MS2-based Jaccard index between two sets of confidently detected peptides. We show evidence that these models can successfully predict the Jaccard index. In addition, we provide a proof of concept that these models can be used to characterize unknown samples without the need to undergo database searching or spectral library matching.

Of all the machine learning models trained, it was observed that there was no single model that obtained the best performance over all datasets. The CNN’s ability to obtain the best performance on the external test set suggests that this model may be more generalizable than the remaining models. The siamese model’s poor performance on the test set, along with its strong performance on both the training and validation set, suggests that the model overfitted to the training data. Our results also suggested that the RF models overtrained as well, given the disproportionate performance of the training set with respect to the test and validation set.

While we demonstrate that our model can be used to characterize samples via unsupervised clustering of the predicted pairwise similarities, future work can be conducted to determine the quality of these clusters using their metadata. Various methods, such as k-means and silhouette scoring, can be used to quantify the cluster quality. However, these methods rely of determining the number of clusters prior to this analysis. Focusing on k-means, another drawback is that the method performs best when the clusters in the graph are spherical, which normally doesn’t occur in cases where data is heavily empirical, such as found in MS-based proteomics data [25]. In addition, it is important to mention that the quality of the unsupervised clustering is dependent on the quality of the associated metadata.

It is known that the generalizability of machine learning models is dependent on the diversity and coverage of training data. It is thereby important to note that one drawback of our training regime is that we only sourced samples from six different organisms, and from PRIDE IDs that had only one organism label. In our results, we found that our models are moderately successful at predicting the Jaccard similarity for runs that originated outside the training dataset. Future work can be done to improve the training dataset to help train more generalized model.

Since our models only take MS1 features as input, they can be used to jointly analyze data collected by different acquisition schemes, such as data-dependent data (DDA) and data-independent data (DIA). This is an advantage over previous methods where DDA and DIA data can not be directly compared due to the different peptide detection strategies. Ultimately, our approach in this study highlights that MS1 scans can be a useful feature in analysis of mass spectrometry runs.

## Availability of data and materials

Code is available at https://github.com/pnnl/deep_scoreMS1. Information on data used in this study can be found in Supplementary File 1 and Supplementary File 2.

## Acknowledgments

Pacific Northwest National Laboratory is a multiprogram national laboratory operated by Battelle Memorial Institute for the United States Department of Energy under contract DE-AC06-76RLO.

## References

[1] Y. Perez-Riverol, A. Csordas, J. Bai, M. Bernal-Llinares, S. Hewapathirana, D. J. Kundu, A. Inuganti, J. Griss, G. Mayer, M. Eisenacher, E. Perez, J. Uszkoreit, J. Pfeuffer, T. Sachsenberg, S. Yilmaz, S. Tiwary, J. Cox, E. Audain, M. Walzer, A. F. Jarnuczak, T. Ternent, A. Brazma, and J. A. Vizcaino. The pride database and related tools and resources in 2019: improving support for quantification data. Nucleic Acids Res, 47(D1):D442–D450, 2019.

[2] M. Choi, J. Carver, C. Chiva, M. Tzouros, T. Huang, T. H. Tsai, B. Pullman, O. M. Bernhardt, R. Huttenhain, G. C. Teo, Y. Perez-Riverol, J. Muntel, M. Muller, S. Goetze, M. Pavlou, E. Verschueren, B. Wollscheid, A. I. Nesvizhskii, L. Reiter, T. Dunkley, E. Sabido, N. Bandeira, and O. Vitek. Massive.quant: a community resource of quantitative mass spectrometry-based proteomics datasets. Nat Methods, 17(10):981–984, 2020.

[3] D. L. Tabb, L. Vega-Montoto, P. A. Rudnick, A. M. Variyath, A. J. Ham, D. M. Bunk, L. E. Kilpatrick, D. D. Billheimer, R. K. Blackman, H. L. Cardasis, S. A. Carr, K. R. Clauser, J. D. Jaffe, K. A. Kowalski, T. A. Neubert, F. E. Regnier, B. Schilling, T. J. Tegeler, M. Wang, P. Wang, J. R. Whiteaker, L. J. Zimmerman, S. J. Fisher, B. W. Gibson, C. R. Kinsinger, M. Mesri, H. Rodriguez, S. E. Stein, P. Tempst, A. G. Paulovich, D. C. Liebler, and C. Spiegelman. Repeatability and reproducibility in proteomic identifications by liquid chromatography-tandem mass spectrometry. J Proteome Res, 9(2):761–76, 2010.

[4] V. Rieder, B. Blank-Landeshammer, M. Stuhr, T. Schell, K. Biß, L. Kollipara, A. Meyer, M. Pfenninger, H. Westphal, A. Sickmann, and J. Rahnenführer. Disms2: A flexible algorithm for direct proteome-wide distance calculation of lc-ms/ms runs. BMC Bioinformatics, 18(1):148, 2017.

[5] M. Palmblad and A. M. Deelder. Molecular phylogenetics by direct comparison of tandem mass spectra. Rapid Commun Mass Spectrom, 26(7):728–732, Apr 2012.

[6] Rob Marissen, Madhushri S. Varunjikar, Jeroen F. J. Laros, Josef D. Rasinger, Benjamin A. Neely, and Magnus Palmblad. comparems2 2.0: An improved software for comparing tandem mass spectrometry datasets. Journal of Proteome Research, 2022.

[7] A. Lin, B. L. Deatherage Kaiser, J. R. Hutchison, J. A. Bilmes, and W. S. Noble. MS1Connect: a mass spectrometry run similarity measure. Bioinformatics, 39(2), Feb 2023.

[8] F. Huber, S. van der Burg, J. J. J. van der Hooft, and L. Ridder. Ms2deepscore: a novel deep learning similarity measure to compare tandem mass spectra. J Cheminform, 13(1):84, 2021.

[9] W. Bittremieux, D. H. May, J. Bilmes, and W. S. Noble. A learned embedding for efficient joint analysis of millions of mass spectra. Nat Methods, 19(6):675–678, 2022.

[10] C. Qin, X. Luo, C. Deng, K. Shu, W. Zhu, J. Griss, H. Hermjakob, M. Bai, and Y. Perez-Riverol. Deep learning embedder method and tool for mass spectra similarity search. J Proteomics, 232:104070, 2021.

[11] Joris Cadow, Matteo Manica, Roland Mathis, Roger R. Reddel, Phillip J. Robinson, Peter J. Wild, Peter G. Hains, Natasha Lucas, Qing Zhong, Tiannan Guo, Ruedi Aebersold, and María Rodríguez Martínez. On the feasibility of deep learning applications using raw mass spectrometry data. Bioinformatics, 37(“1”):i245–i253, 2021.

[12] S. Wang, H. Zhu, H. Zhou, J. Cheng, and H. Yang. MSpectraAI: a powerful platform for deciphering proteome profiling of multi-tumor mass spectrometry data by using deep neural networks. BMC Bioinformatics, 21(1):439, Oct 2020.

[13] William E. Fondrie, Wout Bittremieux, and William S. Noble. ppx: Programmatic access to proteomics data repositories. Journal of Proteome Research, 20(9):4621–4624, 2021.

[14] N. Hulstaert, J. Shofstahl, T. Sachsenberg, M. Walzer, H. Barsnes, L. Martens, and Y. Perez-Riverol. Thermorawfileparser: Modular, scalable, and cross-platform raw file conversion. J Proteome Res, 19(1):537–542, 2020.

[15] H. L. Rost, T. Sachsenberg, S. Aiche, C. Bielow, H. Weisser, F. Aicheler, S. Andreotti, H. C. Ehrlich, P. Gutenbrunner, E. Kenar, X. Liang, S. Nahnsen, L. Nilse, J. Pfeuffer, G. Rosenberger, M. Rurik, U. Schmitt, J. Veit, M. Walzer, D. Wojnar, W. E. Wolski, O. Schilling, J. S. Choudhary, L. Malmstrom, R. Aebersold, K. Reinert, and O. Kohlbacher. Openms: a flexible open-source software platform for mass spectrometry data analysis. Nat Methods, 13(9):741–8, 2016.

[16] J. K. Eng, T. A. Jahan, and M. R. Hoopmann. Comet: an open-source ms/ms sequence database search tool. Proteomics, 13(1):22–4, 2013.

[17] S. McIlwain, K. Tamura, A. Kertesz-Farkas, C. E. Grant, B. Diament, B. Frewen, J. J. Howbert, M. R. Hoopmann, L. Kall, J. K. Eng, M. J. MacCoss, and W. S. Noble. Crux: rapid open source protein tandem mass spectrometry analysis. J Proteome Res, 13(10):4488–91, 2014.

[18] A. Kertesz-Farkas, F. L. Nii Adoquaye Acquaye, K. Bhimani, J. K. Eng, W. E. Fondrie, C. Grant, M. R. Hoopmann, A. Lin, Y. Y. Lu, R. L. Moritz, M. J. MacCoss, and W. S. Noble. The Crux Toolkit for Analysis of Bottom-Up Tandem Mass Spectrometry Proteomics Data. J Proteome Res, 22(2):561–569, Feb 2023.

[19] The UniProt Consortium. UniProt: a worldwide hub of protein knowledge. Nucleic Acids Research, 47(D1):D506–D515, 11 2018.

[20] D. H. May, K. Tamura, and W. S. Noble. Param-medic: A tool for improving ms/ms database search yield by optimizing parameter settings. J Proteome Res, 16(4):1817–1824, 2017.

[21] L. Kall, J. D. Canterbury, J. Weston, W. S. Noble, and M. J. MacCoss. Semi-supervised learning for peptide identification from shotgun proteomics datasets. Nat Methods, 4(11):923–5, 2007.

[22] Günter Klambauer, Thomas Unterthiner, Andreas Mayr, and Sepp Hochreiter. Self-normalizing neural networks. In I. Guyon, U. Von Luxburg, S. Bengio, H. Wallach, R. Fergus, S. Vishwanathan, and R. Garnett, editors, Advances in Neural Information Processing Systems, volume 30. Curran Associates, Inc., 2017.

[23] Yann A. LeCun, Léon Bottou, Genevieve B. Orr, and Klaus-Robert Müller. Efficient BackProp, pages 9–48. Springer Berlin Heidelberg, Berlin, Heidelberg, 2012.

[24] Xavier Glorot and Yoshua Bengio. Understanding the difficulty of training deep feedforward neural networks. In Yee Whye Teh and Mike Titterington, editors, Proceedings of the Thirteenth International Conference on Artificial Intelligence and Statistics, volume 9 of Proceedings of Machine Learning Research, pages 249–256, Chia Laguna Resort, Sardinia, Italy, 13–15 May 2010. PMLR.

[25] A. M. Fahim, A. M. Salem, F. A. Torkey, and M. A. Ramadan. An efficient enhanced k-means clustering algorithm. Journal of Zhejiang University-SCIENCE A, 7(10):1626–1633, 2006.

