## Supplemental Figures for "Predicting the similarity of two mass spectrometry runs using only MS1 data"

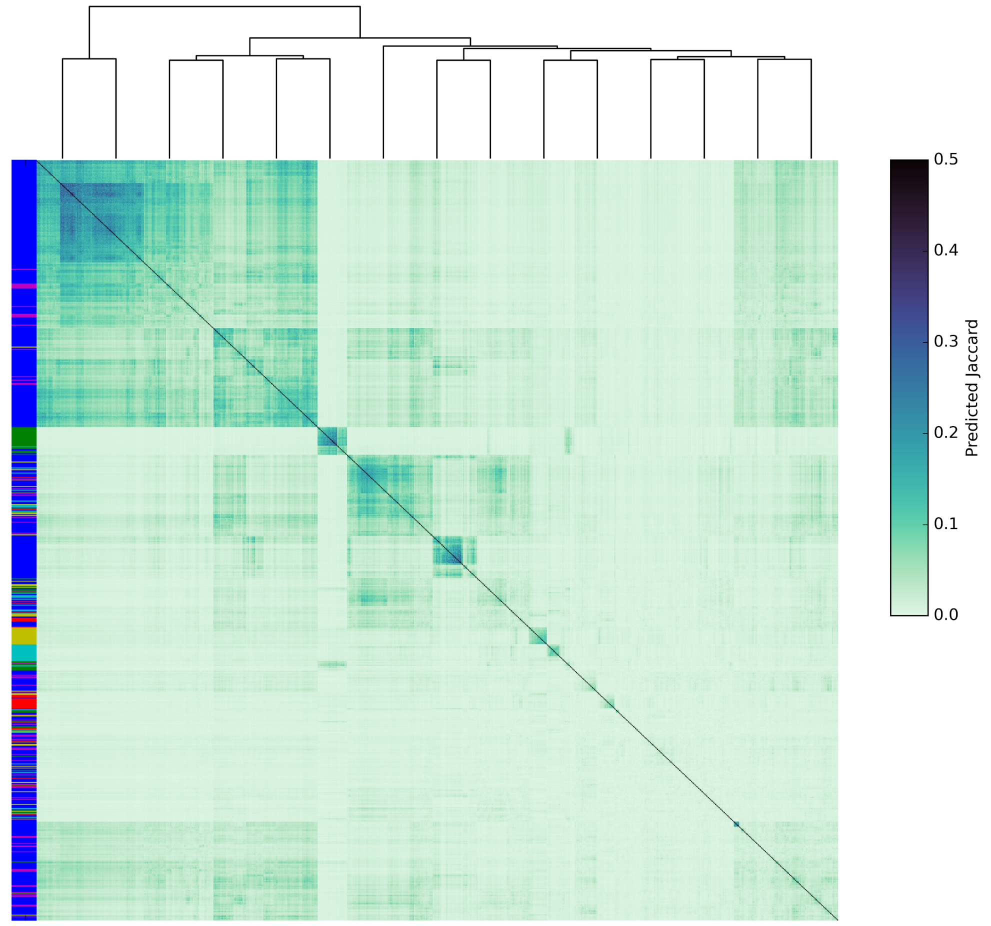


Figure 1: **Hierarchical Clustering of the predicted similarity scores.** A heatmap of the pairwise similarities on the training dataset using the Siamese network. Each cell is colored by the predicted Jaccard index and the denodrogram is created by hierarchical clustering. The bar on the left side of the heatmap is colored by species.

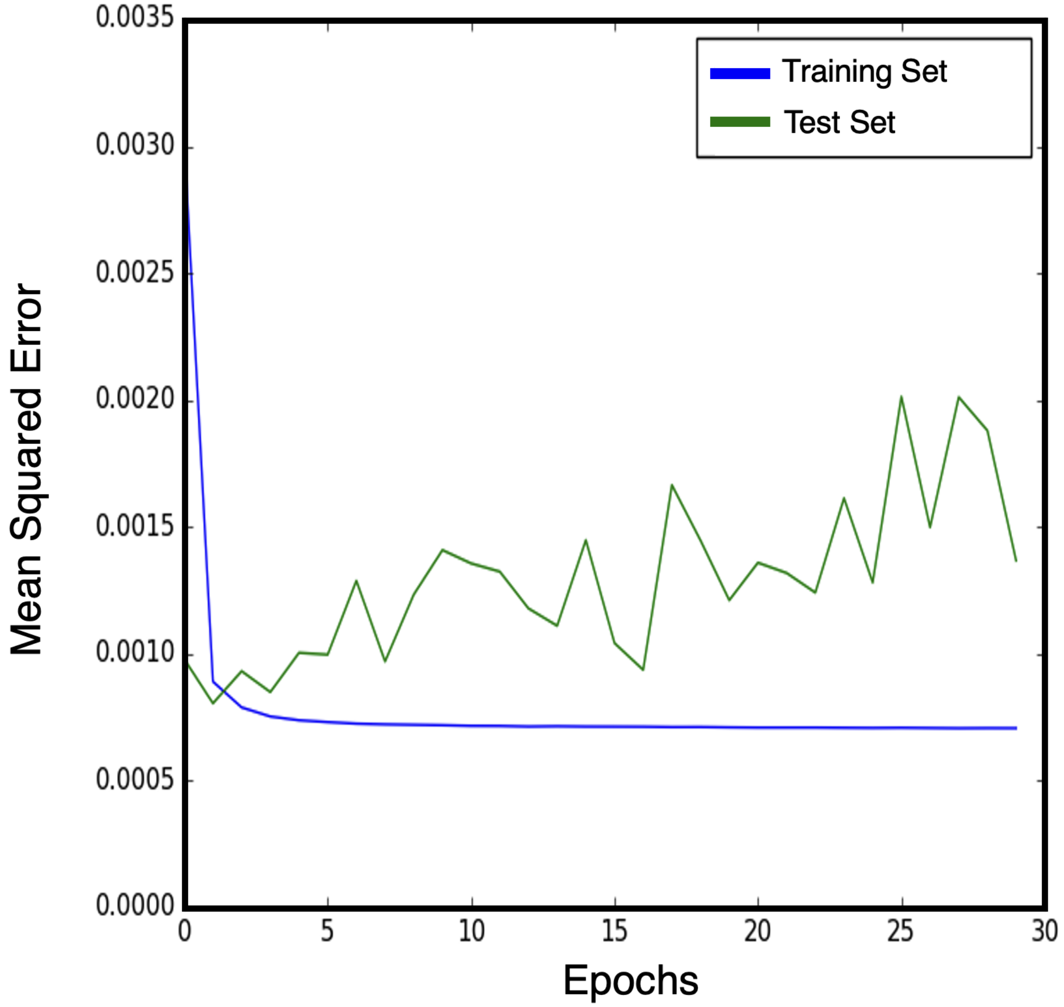


Figure 2: **Loss curve for deep neural network.** A plot of the loss obtained by the deep neural network architecture on the training and test set. The model appears to be overfitted as the test set lose increases as a function of epoch.
